# YTHDF2 is essential for spermatogenesis and fertility through mediating a wave of transcript transition during spermatogonial differentiation

**DOI:** 10.1101/2020.04.30.070235

**Authors:** Xin-Xi Zhao, Zhen Lin, Yong Fan, Yu-Jie Zhang, Fei Li, Tong Hong, Hua Feng, Ming-Han Tong, Ning-Ling Wang, Yan-Ping Kuang, Qi-Feng Lyu

## Abstract

The dynamic and reversible regulation roles of m^6^A modification, and the characterization of m^6^A readers have provided new insights into spermatogenesis at post-transcriptional level. YTHDF2 has been reported to recognize and mediate the m^6^A-containing transcripts decay during the mouse oocyte mature, embryonic stem cell differentiation, neural development, and zebrafish maternal-to-zygotic transition. However, the roles of YTHDF2 in mammalian spermatogenesis are uncertain. Here, we generated germ cell-specific *Ythdf2* mutants (*Ythdf2-vKO*) at a C57BL/6J background, and demonstrated that YTHDF2 was essential for mouse spermatogenesis and fertility. *Ythdf2-vKO* provided oligoasthenoteratozoospermia (OAT) phenotype with increased apoptosis in germ cells. High-throughput RNA-seq of the testis tissue showed the failure of the degradation of a wave of YTHDF2 target mRNA. Interestingly, RNA-seq analysis combined with our previous single-cell transcriptomics data of mouse spermatogenesis pointed out the failure of a wave of transcript transition during the spermatogenesis of *Ythdf2-vKO*, which was confirmed by gene expression analysis of diplotene spermatocytes and round spermatids obtained through fluorescence-activated cell sorting using qPCR. Our study demonstrates the fundamental role of YTHDF2 during mouse spermatogenesis and provides a potential candidate for the diagnosis of male infertility with OAT syndrome.

**Author summary:** Male infertility is becoming a worldwide health problem. Male gamete is generated through spermatogenesis, which is a complicated developmental process with dynamic transcriptome changes. RNA m^6^A modification has been reported as the most prominently internal mRNA modification, which control the tune of gene expression through mRNA splicing, export, translation and decay. RNA m^6^A modification is catalyzed by “writers”, and could be removed by “erasers”. The m^6^A modification enzymes are reported to play important roles during spermatogenesis. Given that the biological function of m^6^A modification are mediated through its “readers”, its readers might be proposed to be involved in the regulation of spermatogenesis. YTHDF2, as a reader of m^6^A modification, has been reported to mediate the m^6^A-containing transcripts decay. To explore the roles of YTHDF2 in spermatogenesis, we used germ cell-specific mutation strategy to knock out the mouse *Ythdf2*. The mutants provided oligoasthenoteratozoospermia phenotype. Our study demonstrate that YTHDF2 is essential for mouse spermatogenesis. YTHDF2 could be a potential candidate for the diagnosis of oligoasthenoteratozoospermia syndrome.

## Introduction

Mammalian spermatogenesis is a highly orchestrated development process typically including three phases: mitosis, meiosis, and spermiogenesis [1]. During mitosis, SSCs undergo self-renew or generate differentiating spermatogonia. In the following meiosis, spermatocytes produce haploid round spermatids through one time replication and two times division. At the third stage, round spermatids undergo a series of morphological and biochemical changes to generate elongated mature spermatozoa [2,3]. These highly complex processes are regulated by epigenetics which will control certain genes to express at different stages. Our previous work and other groups have reported the widely dynamic changes of the transcriptome during mammalian spermatogenesis [4,5]. Moreover, transcription and translation are uncoupling during certain stages of spermiogenesis. The later stage of spermiogenesis is transcriptionally inert [6]. Therefore, the post-transcriptional mechanisms have been one of focus to understand the spermatogenesis [3,7].

Among over 150 types of different RNA modification known to date [8], RNA *N^6^*-methyladenosine (m^6^A), as the most prominently internal mRNA modification in mammalian, has been characterized to tune gene expression through post-transcriptionally regulating m^6^A-containing mRNA metabolism [9,10], including mRNA splicing[11], export[11,12], translation[13] and decay [14,15]. m^6^A has been reported to be involved in extensive biological processes, such as DNA damage response [16], myeloid differentiation [17], embryonic stem cell self-renewal [18], cerebellum development [19], tumorigenicity maintenance [20], T cell homeostasis [21], immune response [22]. RNA m^6^A modification is catalyzed by “writers”, removed by “erasers”, and recognized by its “readers” [9]. The writers are the methyltransferase complex mainly composed of methyltransferase-like 3 (METTL3), methyltransferase-like 14 (METTL14), and Wilms tumor 1-associating protein (WTAP) [9]. We and another group respectively revealed that METTL3/ METTL14 control the SSC fate [13,23]. Furthermore, our group has presented comprehensive m^6^A RNA methylomes during spermatogenesis development stages [13]. The eraser, AlkB homologue 5 (ALKBH5), was reported to modulate the meiotic metaphase-stage spermatocytes and impact the fertility [12]. These reports demonstrated that the dynamic reversible m^6^A modification has fundamental function during mouse spermatogenesis.

The biological functions of m^6^A can be mediated through its readers [9], such as Wnt-driven intestinal stemness amplified by YTHDF1 in m^6^A-dependent manner [24]. The mouse YT521-B homology (YTH) family includes five members, YTH domain family 1 (YTHDF1), YTHDF2, YTHDF3, YTH domain containing 1 (YTHDC1), and YTHDC2 [9,25]. YTHDF1 and YTHDF3 are reported to regulate the translation of their target mRNA through mediating initiation complex loading [26,27]. In contrast, YTHDF2 directs the targets mRNA degradation by recruiting the CCR4-NOT deadenylase complex [15,28]. YTHDC1 regulates mRNA splicing and export [11,29]. YTHDC2 affects mRNA translation and stability [30]. During spermatogenesis, YTHDC1 is essential for spermatogonial development [29]. And YTHDC2 is indispensable for meiotic prophase I through interacting with MEIOC [31]. Besides, YTHDF2 is critical for mouse embryonic stem cell differentiation [32], oocyte maturation and potentiality of early zygotic development [33], neural development [14], and zebrafish maternal-to-zygotic transition [34], through mediating the targets mRNA degradation and turnover. In zebrafish, an average of 70% *ythdf2*^-/-^ progeny arrested at 1-cell stage likely caused by defective sperm [34]. However, *Ythdf2*^-/-^ male mouse showed no defects in spermatogenesis and fertility on a mixed genetic background, although mouse testis displays the highest expression of YTHDF2 [33].

Here, we generated the germline-specific *Ythdf2* knockout mouse at a C57BL/6J background. Our study shows that YTHDF2 is essential for spermatogenesis and fertility. Germline-specific deletion of *Ythdf2*, although had a complete spermatogenesis process, led to oligoasthenoteratozoospermia (OAT), with increased apoptosis in germ cells. RNA-seq of the testis and qPCR analysis of diplotene spermatocytes and round spermatids showed that a wave of transcripts dynamic transition was unsuccessful, which might be caused by a failure of the degradation of YTHDF2 target mRNA timely.

## Results

### Generation of the germ cell-specific deletion of *Ythdf2*

To explore the biological roles of YTHDF2 in mouse spermatogenesis, we first investigated the expression level of YTHDF2 at different development stages. Testes lysates from 3, 7, 14, 21, 28 and 56 days postpartum (dpp) were prepared for western blot. The result showed that YTHDF2 protein was at a low or undetectable level at 3 and 7 dpp stages but dramatically increased from 14 dpp and maintained a high level afterward (Fig 1A). Given that mitosis-to-meiosis transition occurred between 7 dpp and 14 dpp, we speculate that YTHDF2 might play role(s) in process of meiosis and/or spermiogenesis, rather than in mitosis. After clarity of the expression pattern of YTHDF2 in mouse spermatogenesis, we obtained a *Ythdf2*-floxed line in which exon 4 of the *Ythdf2* allele are flanked by *loxP* sites. By crossing with *Vasa-GFPCre* mice, we could specifically delete the fourth exon of *Ythdf2* in male germ cells as early as embryonic day 15 (E15) (S1 Fig). Four genotypes were generated, including *Ythdf2^flox/Δ^ Vasa-GFPCre* (hereafter referred to as *Ythdf2-vKO*), *Ythdf2^flox/Δ^*, *Ythdf2^flox/+^ Vasa-GFPCre, Ythdf2^flox/+^*. The littermate *Ythdf2^flox/+^ Vasa-GFPCre* were used as control (referred as *Ythdf2-CTL*). Both *Ythdf2-vKO* and *Ythdf2-CTL* were healthy and grew normally into adulthood. Agreeing with the Ivanova’s report [33], our immunostaining results also showed that YTHDF2 expressed at each stages of spermatogenesis in control mice, while the signal of YTHDF2 was absent in germ cells of *Ythdf2-vKO* mice, confirming the mutation of *Ythdf2* (Fig 1B). These data showed that *Ythdf2* is dynamically expressed during mouse spermatogenesis, and germ cell-specific *Ythdf2* mutants worked well.

**Fig 1.**
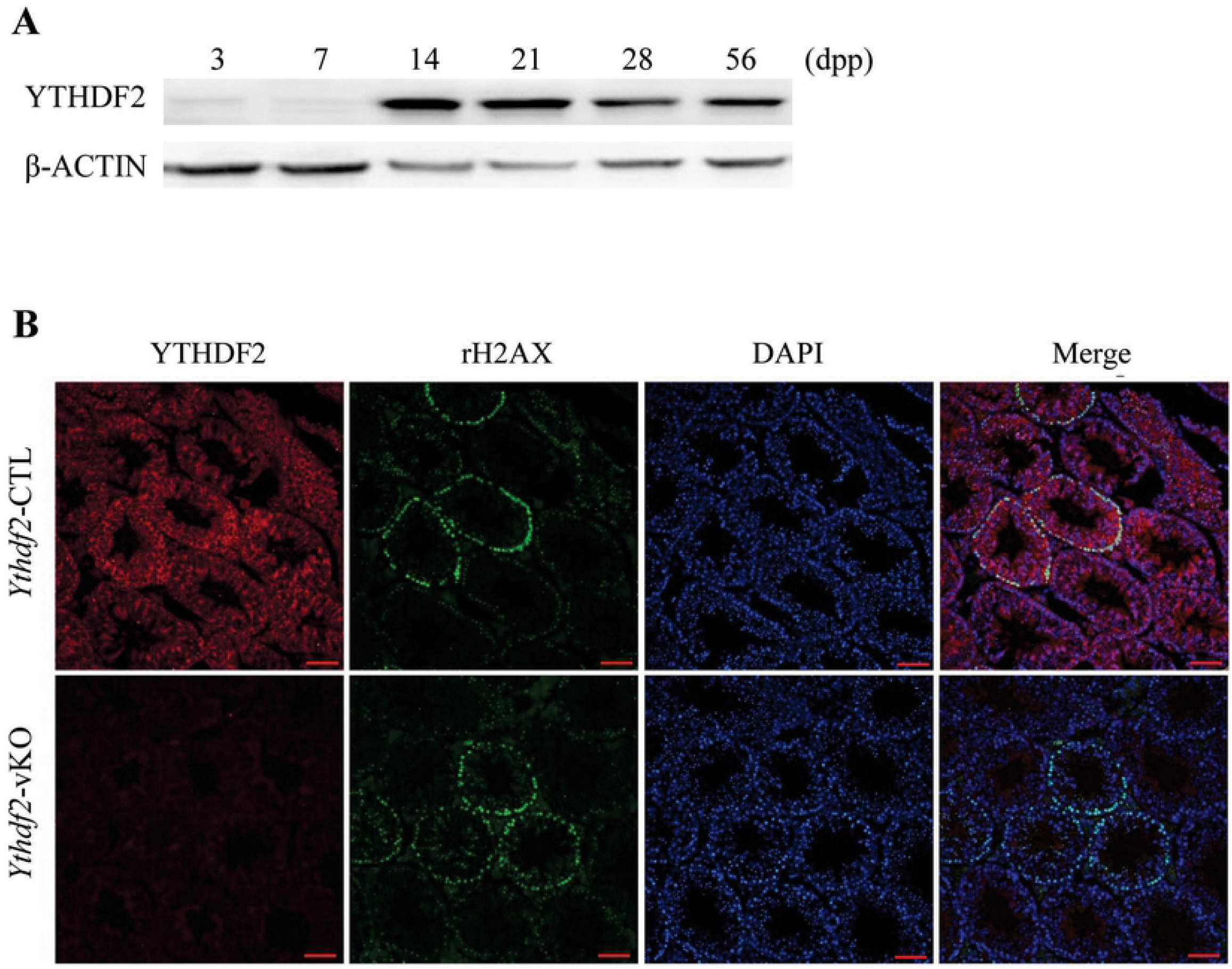
Generation of the germ cell-specific deletion of *Ythdf2*. (A) A representative Western blot of YTHDF2 protein in lysates from wild type testes at different development time. β-ACTIN as loading control. dpp: days postpartum. (B) The efficiency of germ cell-specific deletion of *Ythdf2* was shown through immunofluorescence staining for YTHDF2 (red), the leptotene and zygotene spermatocyte marker γH2AX (green), and DAPI (blue) in sections of 8-week-old control and *Ythdf2-vKO* testes. Scale bar, 40 μm.

### YTHDF2 is required for male fertility and spermatogenesis

To determine the fertility of *Ythdf2*-vKO mice, mating test of *Ythdf2*-vKO male mice with wild-type C57BL/6J (B6) background female mice were carried out. The results showed that *Ythdf2*-vKO could copulate normally, yet were completely sterile (Fig 2A). Although, the testis size of *Ythdf2*-vKO was a little bit smaller than that of *Ythdf2*-CTL (Fig 2B), the weight of testis and body were no significant difference between of them (Fig 2C, D).

**Fig 2.**
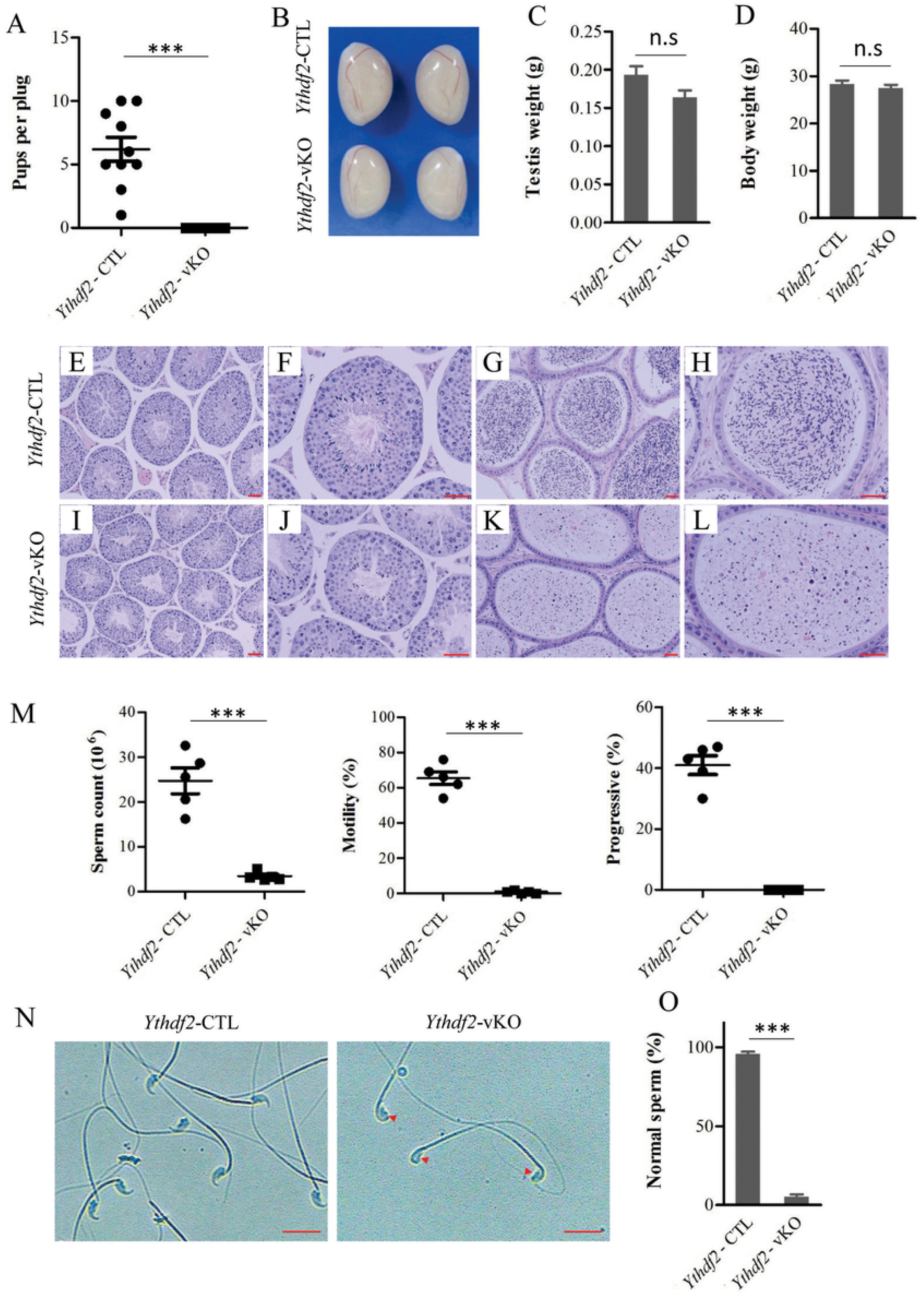
Germ cell-specific *Ythdf2* mutants are characterized with OAT. (A) The infertility of germ cell-specific *Ythdf2* mutants was shown through mating assay. At 2-month-old, control (n=2) and *Ythdf2-vKO* mutant (n=4) were cohabited with two female C57BL/6 mice, respectively. Vaginal plugs were checked and the pups were counted. *** *P* < 0.001, Student’s *t* test. (B) Gross morphology of representative testes from a 2-month-old control and age-matched *Ythdf2*-vKO mice. (C, D) Comparison the testis weight (C) and body weight (D) from 2-month-old controls and age-matched *Ythdf2-vKO* mice. Data are presented as mean ± SEM (n=3 for each genotype). (E-L) H&E staining of testis and epididymis from 2-month-old control (E-H) and age-matched *Ythdf2*-vKO (I-L) mice. Scale bar, 40 μm. (M) The counts, the percentages of motile and progressively motile spermatozoa in cauda epididymides of 2-month-old controls and age-matched *Ythdf2-vKO* mice. Data are presented as mean ± SEM (n=5 for each genotype). *** *P* < 0.001, Student’s *t* test. (N) The morphology of representative sperm from a 2-month-old control and age-matched *Ythdf2*-vKO mice. Red arrow indicates abnormal head sperm. Scale bar, 40 μm. (O) Statistics results of normal sperm from a 2-month-old control and age-matched *Ythdf2-vKO* mice. At least 100 sperm were counted. Data are presented as mean ± SEM (n=3 for each genotype). *** *P* < 0.001, Student’s *t* test.

To characterize the spermatogenesis of *Ythdf2-vKO*, histological analysis showed that spermatogenic cells arranged normally in seminiferous tubules of *Ythdf2-vKO* and *Ythdf2*-CTL adult mice, indicating that *Ythdf2*-vKO possessed a complete spermatogenesis process (Fig 2E, F, I, J). However, the epididymides of *Ythdf2*-vKO contained much less mature spermatozoa than that of *Ythdf2-CTL*, and were filled with a high mount of degenerated germ cells (Fig 2G, H, K, L). Consistent with this, computer-assisted sperm analysis (CASA) showed that the counts, the percentages of motile and progressively motile spermatozoa in *Ythdf2*-vKO cauda epididymides were significantly less than that of *Ythdf2*-CTL (Fig 2M). Besides, more than 90% of the spermatozoa in *Ythdf2*-vKO cauda epididymides were malformed with abnormal heads (Fig 2N, O). Collectively, these data indicated that *Ythdf2*-vKO mutants displayed an OAT phenotype, suggesting that YTHDF2 is required for mouse spermatogenesis and fertility.

### Germ cell-specific *Ythdf2* deletion causes a widely apoptosis during spermatogenesis

Based on the results above, *Ythdf2*-vKO had all kinds of spermatogenic cells in seminiferous tubules and an amount of degenerated germ cells in epididymides, we supposed that apoptosis occurred in seminiferous tubules of *Ythdf2-vKO*. Terminal deoxynucleotidyl transferase-mediated dUTP-biotin nick end labeling (TUNEL) analysis revealed that there were only a few apoptosis signals in seminiferous tubules of *Ythdf2*-CTL, while the seminiferous tubules of *Ythdf2*-vKO contain a significantly increased number of apoptosis cells (Fig 3A-G). Approximately 80% *Ythdf2-vKO* seminiferous tubules had positive apoptosis signals (Fig 3H). These results demonstrated that a widely apoptosis happened in seminiferous tubules of *Ythdf2*-vKO, which might explain the degenerated germ cells in epididymides.

**Fig 3.**
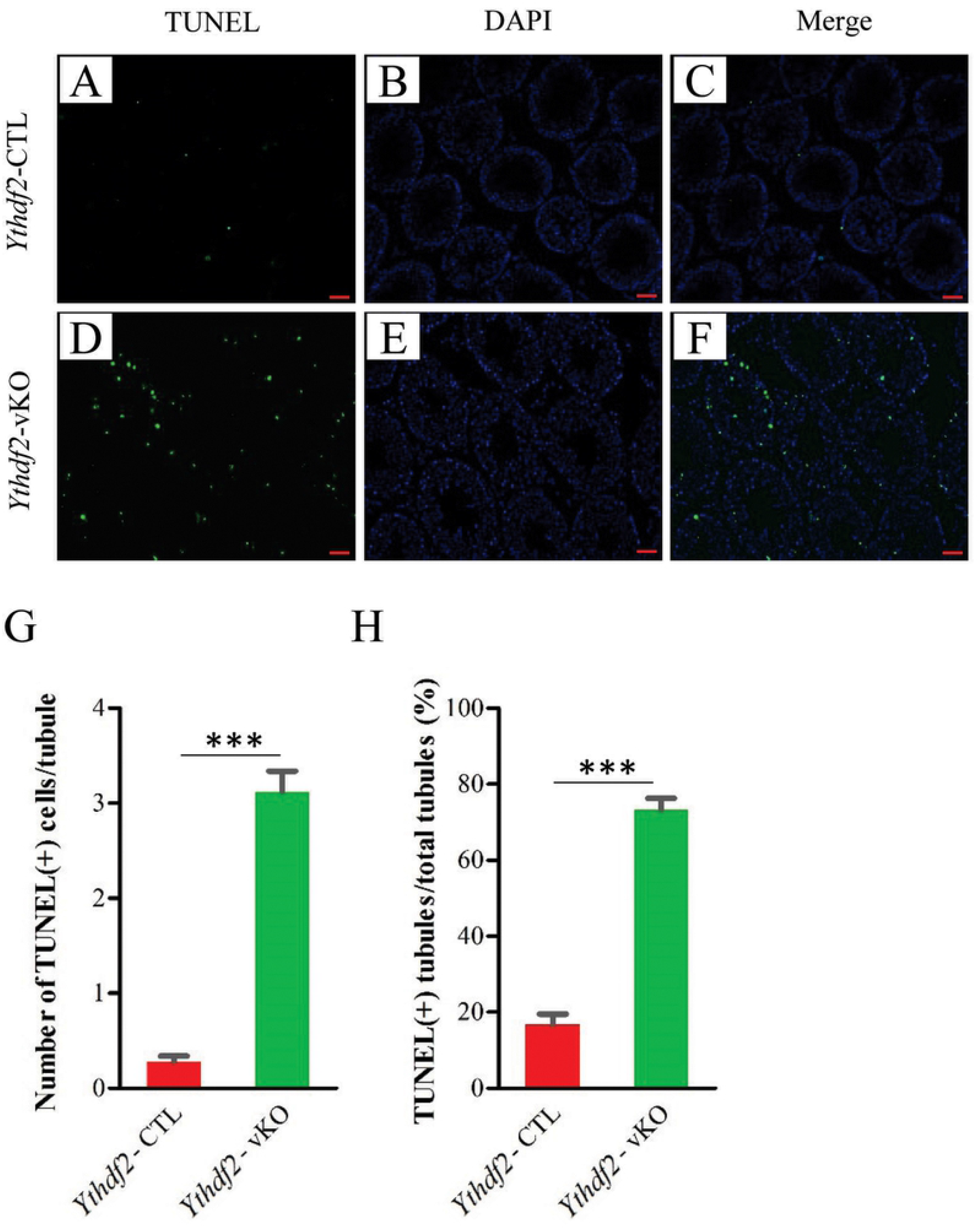
Germ cell-specific *Ythdf2* deletion causes a severe apoptosis during spermatogenesis. (A-F) Representative of TUNEL staining of testes from 2-month-old controls (A-C) and age-matched *Ythdf2-vKO* (D-F) mice. The staining of apoptotic cells with TUNEL (green) and of the nucleus with DAPI (blue). Scale bar, 40 μm. (G) Quantification of TUNEL-positive cells in seminiferous tubules from 2-month-old controls and age-matched *Ythdf2-vKO* mice. Data are presented as mean ± SEM (n=3 for each genotype; tubules examined: 185 of control and 205 of *Ythdf2-vKO* mice). *** *P* < 0.001, Student’s *t* test. (H) Comparison of TUNEL-positive tubules from 2-month-old controls and age-matched *Ythdf2-vKO* mice. Data are presented as mean ± SEM (n=3 for each genotype; tubules examined: 185 of control and 205 of *Ythdf2-vKO* mice). *** *P* < 0.001, Student’s *t* test.

### YTHDF2 regulates dynamic mRNA dosage during spermatogenesis

Given the reported molecular role of YTHDF2 in mRNA stability, we conducted high-throughput RNA sequencing (RNA-seq) on testes from *Ythdf2*-vKO and *Ythdf2*-CTL adult mice to investigate the biological mechanism underlying the defects of *Ythdf2*-vKO spermatogenesis. Compared with *Ythdf2*-CTL, we identified 266 genes (196 up and 70 down) deregulated in *Ythdf2-vKO* testes (*P* < 0.05, fold change > 1.5) (Fig 4A, S1A table). Our qPCR data confirmed the RNA-seq analysis (Fig 4B). A broad of biological processes enriched for the upregulated genes in *Ythdf2*-vKO mice (Fig 4C, S1B table). According to our previously reported m^6^A mRNA methylomes of mouse spermaogenic cells [13], 66% of the upregulated genes (129 genes) in *Ythdf2-vKO* testes were modified with m^6^A (Fig 4D). The distribution of m^6^A of the 129 genes in wild-type transcriptome were mainly at 3UTR, Stop Codon, and CDS (74% in total) (Fig 4D). The deregulated genes chiefly toward upregulation in *Ythdf2*-vKO testes (Fig 4A), and most of which were modified with m^6^A (Fig 4D), are naturally consequent on the deletion of *Ythdf2* that involves mRNA decay.

**Fig 4.**
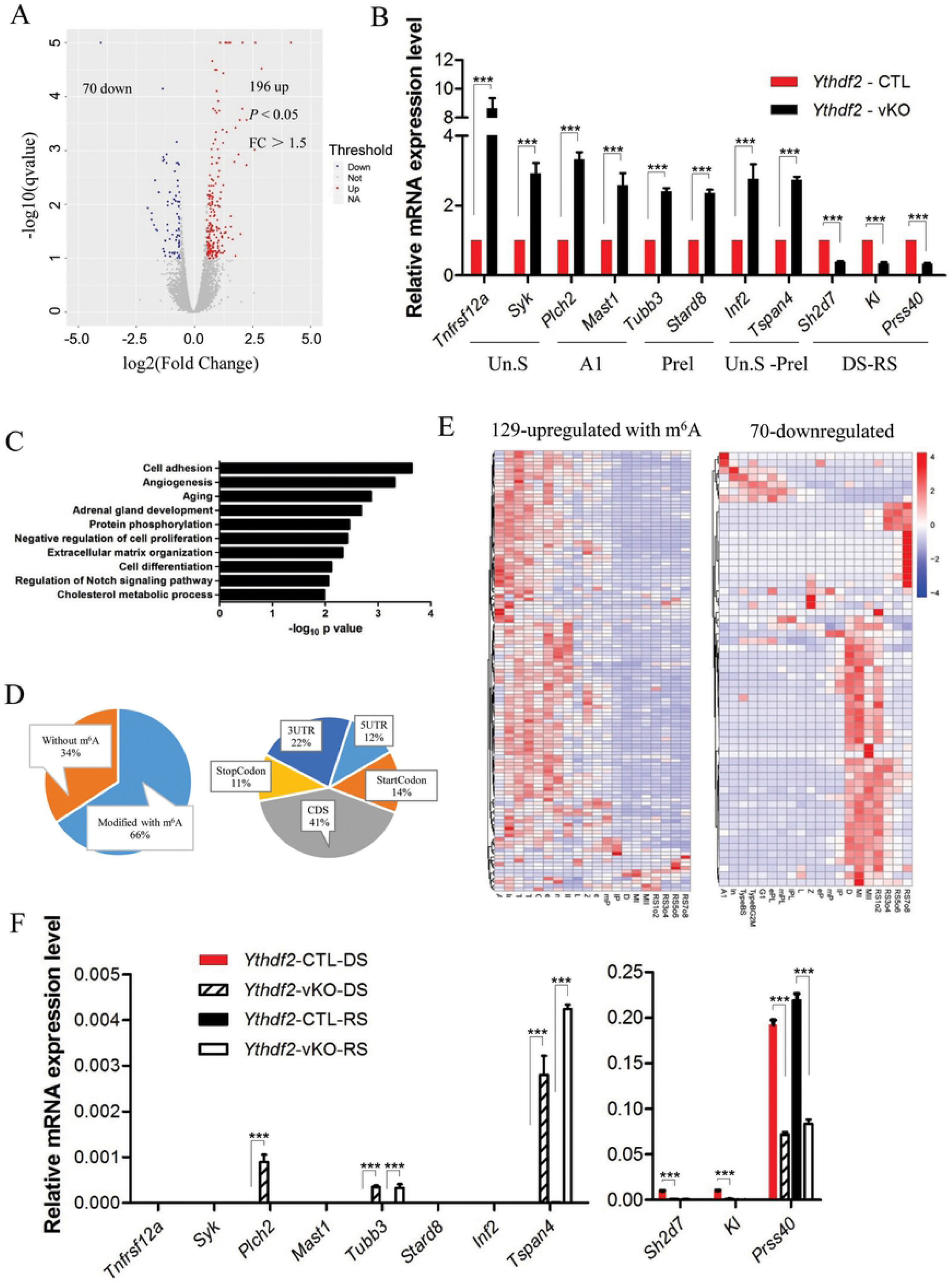
Germ cell-specific *Ythdf2* deletion alters dynamic expression pattern of genes during spermatogenesis. (A) Scatter plot showing the fold change of transcripts of testis tissue between 2-month-old controls and age-matched *Ythdf2-vKO* mice. Significantly downregulated and upregulated (p < 0.05) genes with a fold change greater than 1.5 are shown in blue and red, respectively. Analysis was done on two biological replicates. (B) qPCR validation of the deregulated genes in *Ythdf2-vKO* spermatogenesis. Un.S, undifferentiated spermatogonia; A1, type A1 spermatogonia; Prel, preleptotene spermatocytes; DS, diplotene spermatocytes; RS, round spermatids. Data are presented as mean ± SEM (n=3 for each genotype). *** *P* < 0.001, Student’s *t* test. (C) Gene ontology enrichment analysis for the upregulated genes in *Ythdf2*-vKO mice. The top ten most enriched biological processes are shown. (D) The ratio of transcripts modified with m^6^A among the upregulated genes in (a) and the distribution of m^6^A sites were analyzed according to our previously reported m^6^A mRNA methylomes of mouse spermaogenic cells. (E) Heatmap analysis of the wild-type expression level of the upregulated and downregulated genes in *Ythdf2*-vKO mice based on our previous published single-cell transcriptomics analyses of mouse spermatogenesis. A1, type A1 spermatogonia; In, intermediate spermatogonia; TypeBS, S phase type B spermatogonia; TypeBG2M, G2/M phase type B spermatogonia; G1, G1 phase preleptotene; ePL, early S phase preleptotene; mPL, middle S phase preleptotene; lPL, late S phase preleptotene; L, leptotene; Z, zygotene; eP, early pachytene; mP, middle pachytene; lP, late pachytene; D, diplotene; MI, metaphase I; MII, metaphase II; RS1o2, steps 1–2 spermatids; RS3o4, steps 3–4 spermatids; RS5o6, steps 5–6 spermatids; RS7o8, steps 7–8 spermatids. (F) The changes of dynamic expression pattern of genes during spermatogenesis of germ cell-specific *Ythdf2* mutants were confirmed using qPCR analysis of two stages of spermatogenic cells (diplotene spermatocytes and round spermatids) collected through FACS. DS, diplotene spermatocytes; RS, round spermatids. Data are presented as mean ± SEM (n=3 for each genotype). *** *P* < 0.001, Student’s *t* test.

Notably, the 129 upregulated genes with m^6^A modification in *Ythdf2*-vKO testes were mainly distributed at early stages of spermatogenesis in our previously reported m^6^A mRNA methylomes of mouse spermaogenic cells [13] (S2 Fig). To confirm this trendency, we further analyzed the precision distribution at single-cell level of the 129 upregulated genes using our published single-cell transcriptomics data of mouse spermatogenesis [5] (Fig 4E). These two analysis consistently indicate that the 129 upregulated genes with m^6^A modification in *Ythdf2*-vKO testes mainly express at early stages of wild-type spermatogenesis. While the 70 down regulated genes in *Ythdf2*-vKO testes mainly exist at later stages of wild-type spermatogenesis, specially begin to express at diplotene spermatocytes of wild-type mice (Fig 4E). To verify the expression changes of differentially expressed genes in *Ythdf2-vKO*, we performed qPCR analysis on two types of germ cells, diplotene spermatocytes and round spermatids collected through fluorescence-activated cell sorting (FACS) from *Ythdf2-vKO* and *Ythdf2-CTL* testes. Our data showed that 3 of the 8 genes expressed at early stages during spermatogenesis had a significantly high level in diplotene spermatocytes of *Ythdf2*-vKO than that of *Ythdf2*-CTL, and the expression of 2 in these 3 genes were still significantly higher in round spermatids stage of *Ythdf2-vKO*. All of the level of 3 genes that expressed at diplotene spermatocytes backward in wild-type mouse decreased significantly in diplotene spermatocytes of *Ythdf2-vKO* than that of *Ythdf2-CTL*, and 1 in 3 genes also had a lower level in round spermatids stage of *Ythdf2*-vKO (Fig 4F).

Collectively, these results showed that YTHDF2 is essential for regulation of a wave of transcriptome dynamic transition during mice spermatogenesis perhaps through removal of useless transcripts which are modified with m^6^A.

## Discussion

Emerging evidences have shown the dynamic and reversible regulation roles of m^6^A modification in mammalian spermatogenesis, particularly the characterization of m^6^A readers has provided new insights into post-transcriptional mechanisms in spermatogenesis [3,29,31]. However, the biological roles of YTHDF2 in mammalian spermatogenesis are uncertain. In this study, we generated a germ-cell specific *Ythdf2* knockout mouse at a C57BL/6J background (Fig 1). Our results demonstrate that YTHDF2 is critical for spermatogenesis and fertility (Fig 2). Deletion of *Ythdf2* led to OAT and increased apoptosis in germ cells (Fig 3). High-throughput RNA-seq of the testis tissue and qPCR analysis of diplotene spermatocytes and round spermatids confirmed that the degradation of a wave of YTHDF2 target mRNA failed, and the expression of following subset genes were affected (Fig 4E, F).

Based on our results of the dynamic expression of YTHDF2 at developmental stages (Fig 1A) and the reported special highest expression level of YTHDF2 in mouse testis tissue [33], and the reported clue of sperm defects in zebrafish [34], we proposed that YTHDF2 play critical roles during mammalian spermatogenesis. Disaccording with the report on a mixed genetic background [33], our study show that YTHDF2 is required for mouse spermatogenesis and fertility at C57BL/6J background (Figs 2 and 3). Furthermore, the phenotypes of *Ythdf2-vKO* are very similar with *Alkbh5*^-/-^ mice, which also provided the following characters, such as OAT, increasing apoptosis, and infertility [12]. Given the existence of each types of germ cells in seminiferous tubule of *Ythdf2-vKO*, the spermatogenic maturation and process of spermiogenesis of *Ythdf2-vKO* requires further study.

The high complex of spermatogenesis manifests the dynamic transcriptome and m^6^A RNA methylomes of different stage spermatogenic cells to timely-tune mRNA translation and decay at post-transcriptional level [3,13]. The install of m^6^A on transcripts could mark them for recognition by readers. According with the regulation role of YTHDF2 on mRNA stability, deletion of YTHDF2 led to 196 transcripts upregulated, 129 of which were marked with m^6^A (Fig 4A, D). Likely YTHDC2 [31], only a small part of transcripts among the large number of genes marked with m^6^A was targeted by different reader to accelerate decay. This implies the existence of more readers or more coordinating mechanisms of m^6^A-containing transcripts at post-transcriptional level, which are worthy for investigation in future.

Interestingly, we found that there was a wave of transcripts transition failure (Fig 4E) through mapping the deregulated transcripts *Ythdf2*-vKO on our previously reported single-cell transcriptomics of mouse spermatogenesis [5]. Specially, the expression of a testis-specific gene, *protease, serine 40* (*Prss40*) [35], was downregulated in *Ythdf2-vKO* mutant testes (Fig 4B). Besides, the absence of a serine protease inhibitor, *Spink2*, led to OAT in heterozygotes and azoospermia in homozygotes [36]. The causal relationship of the downregulation of *prss40* and OAT is worthy for further study. Given the amount of RNA from germ cells limiting the performing RNA-seq, we confirmed the transcripts transition failure in *Ythdf2*-vKO using qPCR analysis of diplotene spermatocytes and round spermatids obtained through FACS (Fig 4F). Our data and other studies in embryonic stem cell [32], oocyte mature [33], neural development [14], and zebrafish maternal-to-zygotic transition [34], together support the hypothesis that YTHDF2 plays a crucial role in different development transiting stage through recognition and clearance the no longer necessary targets, giving road to express genes required in following development stages.

In summary, our study show that YTHDF2 is required for mouse spermatogenesis and fertility at C57BL/6J background. YTHDF2 mediates a wave of transcripts transition, the failure of which potentially is the cause of apoptosis of germ cells in seminiferous tubule of *Ythdf2-vKO*. The explanation for the deregulation of the subset of gene expression and the arising of apoptosis in *Ythdf2*-vKO testis needs more detailed study in further.

## Materials and methods

### Mice

The *Ythdf2* knockout first mice were obtained from Model Animal Research Center of Nanjing University. In the knockout first allele, a promoter-driven cassette (including *LacZ* and *neo* genes) flanked by FRT sites are inserted between exon 3 and exon 4 whereas the exon 4 of *Ythdf2* are flanked by *LoxP* sites. We converted the knockout first allele to a conditional allele by crossing *Ythdf2* knockout first mice with *Flp* deleter mice. The resulting floxed *Ythdf2* mice were crossed with *Vasa-GFPCre* knock-in mice (generated by Shanghai Biomodel Organism Co., Ltd.) to allow specific knockout of *Ythdf2* in germ cells. All mice were kept on the C57BL/6J background. The primers used for genotyping were listed in S2 table. *Ythdf2^flox/Δ^ Vasa-GFPCre* (*Ythdf2-vKO*) was identified for this study, and *Ythdf2^flox/+^ Vasa-GFPCre* (*Ythdf2*-CTL) as control. All animal experiments were conducted in accordance with the guidelines of the Animal Care and Use Committee at Shanghai Institute of Biochemistry and Cell Biology, Chinese Academy of Sciences.

### Isolation of spermatogenic cells

After the two-step enzymatic digestion and Hoechst 33342 staining as described previously [13], two types of spermatogenic cells from *Ythdf2-vKO* and control, diplotene spermatocytes and round spermatids, were isolated using FACS (Becton Dickinson).

### Total RNA isolation and quantitative RT-PCR

Total RNA was isolated from testis tissue or spermatogenic cells using TRIzol Reagent (15596018, Ambion/Life Technologies) according to the manufacturer’s instructions. RNA quality was assessed by a NanoDrop 2000, and cDNA was prepared by using a commercial kit (FSQ-301, Toyobo). RT-qPCR was performed on an Eppendorf Mastercycler PCR System using SYBR Green Mix (QPK-201, Toyobo). The primers were designed using Primer3 (Version. 0.4.0) and synthesized at Generay Biotech (Shanghai, China). The relative expression level was normalized to *β-actin*.

### Western blotting

For Western blotting analysis, the testis tissues were homogenized using TGrinder (OSE-Y10-Plus, Tiangen Biotech), and lysed in 1× SDS loading buffer by boiling for 10 min. 10% SDS-PAGE was used to resolve the proteins, which were then transferred to a nitrocellulose membrane. After blocked with 5% nonfat milk, the membranes were incubated with anti-YTHDF2 (1:5000; Proteintech, 24744-1-AP) overnight at 4 °C. Goat anti-rabbit IgG, HRP (1:5000, Thermo Fisher, 31460) was used as secondary antibody. Anti-β-ACTIN (1:3000, AB2001, Abways) as control. SuperSignal™ West Pico PLUS Chemiluminescent Substrate (Thermo Scientific, 34577) was used to detect the antibody binding.

### Histology and immunofluorescence staining

For histology, testes from *Ythdf2-vKO* and control were fixed in Bouin’s solution, embedded in paraffin and sectioned. Hematoxylin and Eosin (H&E) was carried out after Sections deparaffinized and rehydrated. For immunofluorescence analysis, testes were fixed in 4% paraformaldehyde (PFA), embedded in paraffin and sectioned. After deparaffinization and rehydration, sections were boiled in 10 mM sodium citrate buffer (pH 6.0) for 15 min, washed in PBS with 0.1% Triton X-100. The sections were then blocked with blocking buffer (10% donkey serum and 0.1% Triton X-100 in PBS) for 60 min, and then incubated with the primary antibodies overnight at 4 °C. The following primary antibodies were used: anti-YTHDF2 (1:100; Proteintech, 24744-1-AP) and mouse anti-γH2AX (1:500; Millipore, 05-636).

### TUNEL analysis

Testes from *Ythdf2*-vKO and control were fixed in 4% PFA, embedded in paraffin and sectioned. After deparaffinization and rehydration, sections were boiled in 10 mM sodium citrate buffer (pH 6.0) for 1 min, and then added ddH_2_O and cooled to room temperature. The sections were blocked with blocking buffer (10% donkey serum and 0.1% Triton X-100 in PBS) for 30 min. The TUNEL staining was performed using the TUNEL BrightRed Apoptosis Detection Kit (Vazyme, A112-02) according to the manufacturer’s instructions. The sections were then washed in PBST three times, and mounted on slices with Mounting Medium with DAPI-Aqueous, Fluoroshield (Abcom, ab104139). Images were observed using fluorescence microscope (Zeiss Axio Scope A1).

### CASA

Cauda epididymides were separated from control and Ythdf2-vKO and then minced in pre-warmed (37°C) Tyrode’s Buffer (Sigma-Aldrich, USA). After 15 minutes at 37 °C, tissue was removed, the sperm suspension assessment including the count, the percentages of motile and progressively motile spermatozoa were determined by CASA (Hamilton Thorne, USA).

### RNA-seq

The testes from control and Ythdf2-vKO were homogenized using TGrinder (OSE-Y10-Plus, Tiangen Biotech) and prepared for RNA extraction. RNA quality was assessed using a bioanalyzer (Agilent Technologies). RNA-seq libraries were prepared according to the protocol for use with NEBNext rRNA Depletion Kit (Human/Mouse/Rat) (NEB #E6310) and NEBNext Ultra II Directional RNA Library Prep Kit for Illumina (NEB #E7760). Then were pooled for deep sequencing by using Illumina Hiseq Xten (2 × 150) platforms at the CAS-MPG Partner Institute for Computational Biology Omics Core, Shanghai, China. Raw read qualities were evaluated with FastQC (http://www.bioinformatics.babraham.ac.uk/projects/fastqc/,v0.11.5). Adaptor sequences and read sequences on both ends with Phred quality scores below 30 were trimmed. Trimmed reads were then mapped with the Hisat2 algorithm (Hisat2 v2.1.0) to target sequences. Gene expression levels were quantified by the software package HTSeq (v0.6.1p1). Differentially expressed genes were generated by asking for a log2 (-fold change) > 1.5 with *P* < 0.05. The significant genes were subsequently analyzed for enrichment of biological themes using the DAVID 6.8 bioinformatics platform (https://david.ncifcrf.gov/). Data from this experiment has been deposited in the GEO (GSE147574) and NODE (OEP000805, https://www.biosino.org/node/project/detail/OEP000805) database.

### Statistics

Student’s *t* test was used for statistical analyses by using GraphPad Prism software (Version 5.01). A *P* < 0.05 indicated a statistically significant difference between groups. The data are presented as the mean ± SEM.

## Supporting Information Legends

**S1 Fig. Strategy for the generation of the germ cell-specific deletion of *Ythdf2*.**

A *Ythdf2-floxed* line was generated, in which exon 4 of the *Ythdf2* if flanked by *loxP* sites. The germ cell-specific *Ythdf2* mutants were obtained by crossing with *Vasa - GFPCre* mice.

**S2 Fig. The wild-type expression stages of the upregulated genes with m^6^A modificaiton in *Ythdf2-vKO*.**

Mapping the 196 upregulated genes in our previous data of mouse spermatogenesis with five stages and counting the number of genes in each stage.

**S1 Table. Differentially expressed genes in the testis tissue of Ythdf2-vKO and the GO analysis.**

**S2 Table. The sequences of primers.**

